# HYPERTRIGLICERIDEMIC WAIST, PHYSICAL ACTIVITY AND CARDIOVASCULAR RISK FACTORS IN SCHOOLCHILDREN

**DOI:** 10.1101/2020.06.05.136101

**Authors:** Flávio Ricardo Guilherme, Stevan Ricardo dos Santos, Rodrigo Garcia Fiorillo, Wilson Rinaldi

**Author notes:** **Contributors:** F.R Guilherme participated in all stages of the study, including the conception, data analysis and interpretation, and writing and final approval of the article. S.R dos Santos and R.G Fiorillo participated in the data analysis and interpretation and writing and final approval of the article. W. Rinaldi participated in the study’s conception and development and critical revision of the article.

## Abstract

The present study aimed to analyze the prevalence of the Hypertriglyceridemic Waist (HTW) and its rates with levels of physical activity, nutritional status and lipid profile of students from public schools. A sample consisted of 448 schoolchildren between 10 and 18 years old, who are assessed by body mass, height, BMI (waist circumference), triglycerides, total cholesterol, HDL-C, LDL-C, non-HDL cholesterol, fasting blood glucose and weekly physical activity time. The survey results showed a high prevalence of the HTW phenotype in schoolchildren (n = 125; 27.9%). The group diagnosed with phenotype has higher body mass, BMI, total cholesterol, LDL-C and non-HDL cholesterol and a lower level of HDL-C about the group without a diagnosis of the phenotype. However, for fasting blood glucose measurements and time of physical activity, the groups are no different. An association analysis using logistic regression showed the HTW phenotype associated with sex, nutritional status, and total cholesterol, where boys presented 2.0 (95% CI 1.3 - 3.2); obese 6.2 (95% CI 2.7 - 17.2) and cholesterol levels above 150 mg / dL 3.5 (95% CI 2.1 - 5.7) times more likely to have the phenotype. In this way, the present research showed a high prevalence of HTW in schoolchildren from public schools, with boys, obese and schoolchildren with total cholesterol levels, the individuals most likely to have the phenotype. However, it is worth emphasizing the importance of monitoring these variables throughout the year in all courses, given the period of strong, biological, behavioral and psychological changes, as they can quickly change the values of the analyzed variables.

## Introduction

The term hypertriglyceridemic waist (HTW) started to be used around the 2000s when Lemieux et al., (2000) ^1^ investigated the hypothesis that variables considered simple such as waist circumference (WC) and plasma triglyceride levels (TG) fasting could be used as useful tools in the identification of risk factors for the development of cardiovascular diseases.

Currently, the HTW phenotype is defined as the joint presence of elevated levels of CC and TG, ^2^, and those diagnosed in turn, have a higher chance of having increased blood glucose and systemic arterial hypertension, being more prone to the development of the metabolic syndrome and pathologies classified as risk factors for cardiovascular diseases ^2^.

To prevent the HTW phenotype and the risk factors associated with the phenotype in children and adolescents, regular physical activity is recommended^3^, because, in addition to the preventive characters in these age groups, good levels of physical activity provide greater health benefits, presenting better cardiorespiratory and muscular fitness, bone health and maintenance of adequate body weight ^4^.

Despite the association of HTW with body adiposity ^3^ and consequently the increased cardiometabolic risk in children and adolescents ^5^, few studies have verified the prevalence and association of this phenotype with the level of physical activity and cardiovascular risk factors in public school students in Brazil. Because of the above, the present study aimed to analyze the prevalence of the HTW phenotype and its associations with the level of physical activity, cardiovascular risk factors in students from public schools in Paranavaí, Brazil.

## Methods

Research with a cross-sectional design, sample composed of students of both sexes aged 10 to 18 years, from the public network of the morning period of the city of Paranavaí, Paraná State, Brazil. According to data from the city’s Regional Education Center, 3,483 students were enrolled in the city’s eight schools. The classes were selected by intentional sampling, structured in two stages: 1) Invitation to students from all classes and explanation of study; 2) delivery of the Consent Term for signature by parents or legal guardians.

The sample calculation resulted from the total number of the population (n=3.483); a prevalence of 20.7% of HTW in schoolchildren in the same city in 2014 ^6^ confidence level equal to 95%; and sampling error of 4%. Based on these parameters, data from 354 students were required. 10% of this value was added to the sample, predicting eventual losses and refusals (354 + 70 = 389), resulting in the collection of data from 389 adolescents for selection of the final sample. Assessments were made only for those students who delivered the informed consent form signed by parents and / or guardians, totaling 448 students.

Collections were performed during school hours by trained evaluators who used previously calibrated equipment. Height was measured with a wall stadiometer (Wisoâ, Brazil) and body mass on a digital scale (G-Tech) with a maximum capacity of 150kg. During the evaluations the students wore only a school uniform, with no objects in their pockets. The Body Mass Index (BMI) was used to classify the nutritional status in eutrophic, overweight and obese individuals ^7^.

Central obesity was assessed using waist circumference, obtained using a flexible and inextensible measuring tape (Gulick, Brazil), with a resolution of 0.1 cm applied above the iliac crests. For classification, the cut-off point ≥ P75 was used for all ethnicities ^8^.

Biochemical tests were requested only for adolescents who had central obesity and accepted to perform the collection (n = 448). For this purpose, a 10 ml sample of venous blood was collected in the anterior cubital vein after a fasting period of at least 10 hours, between 8 am and 9:30 am in the schools themselves. The samples were properly collected, centrifuged at 1500g for 15 minutes at 4° C and analyzed on the same day in a clinical analysis laboratory in the city. Serum levels of total cholesterol, high-density lipoprotein cholesterol (HDL-C), non-HDL cholesterol, low-density lipoprotein cholesterol (LDL-C), triglycerides and fasting blood glucose (GJ) were determined by enzymatic and colorimetric methods with Gold Analyzes kit, according to the manufacturer’s specifications. The values of total cholesterol≥150 mg / dL, LDL≥100 mg / dL, HDL <45 mg / dL, non-HDL ≥123 mg / dL, triglycerides≥100 mg / dL and fasting glucose ≥100 mg / dL were considered inadequate ^(3,9)^.

The HTW phenotype was defined by the simultaneous presence of abdominal obesity (P75) FERNÁNDEZ et al., (2004) ^8^ and high serum triglyceride levels (≥100 mg/dl)^10^.

The level of physical activity was assessed using the questionnaire for adolescents^11^,adapted from the Self-Administered Physical Activity Checklist^12^.The questionnaire contains a list of 24 activities of moderate/vigorous intensity, with the option for the subject to add two more in the case of activities they performed, but are not on the list. For each activity listed, the student can enter the frequency (days/week) and duration (hours/min/day) of physical activities practiced in the last seven days, which resulted in weekly physical activity time. The cut-off point for inadequate physical activity level was <300 minutes/week ^13^.

In the statistical analysis, the Kolmogorov Smirnov test was used to identify the normality of the data. Descriptive analysis was performed by means and standard deviation. Mann Whitney U test was used to compare the groups, due to the absence of parametric data distribution. The Chi-Square test was used to verify the difference in the proportion of HTW (dependent variable), according to the categories of independent variables, and for 2×2 contingency tables, Yates Continuity Correction was performed. Multivariate logistic regression was used to determine the odds ratio or Odds Ratio (OR) and the respective confidence intervals (95%), to analyze the association of HTW with the independent variables. The criterion for including the independent variables in the multivariate model was an association level of p≤0.20 with the dependent variable by the Chi-square test, and permanence in the model if there was an association (p≤0.05) in the multivariate analysis. The analyzes were performed using the Statistical Package for Social Science (SPSS), version 20.0, considering p≤0.05.

This study was approved by the Research Ethics Committee of the State University of Maringá, under opinion number 1.453.730, according to the Declaration of Helsinki.

## Results

Table 1 compares the means between the groups without a diagnosis of and without the presence of the HTW phenotype. The data show the similarity between groups for the means of age, height, time of physical activity and fasting blood glucose. However, for body mass, BMI, total cholesterol, LDL-C and non-HDL cholesterol, the group diagnosed with HTW (HTW +) had higher means (p <0.001) than the group without a diagnosis (HTW−), and lower for HDL-C (p <0.001).

**Table 1.** Time of physical activity and health risk factors in schoolchildren with and without a diagnosis of HTW.

| Variables | Mean ± SD | | p-value |
| --- | --- | --- | --- |
|  | HTW- (n=323) | HTW+ (n=125) |  |
| Age (years) | 13.58 ± 1.87 | 13.94 ± 2.03 | 0.119 |
| PA Time (min) | 732.9 ± 986.7 | 645.2 ± 716.2 | 0.379 |
| Body Weight (kg) | 67.6 ± 13.82 | 74.65 ± 16.76 | < 0.001* |
| Height (cm) | 1.62 ± 0.10 | 1.64 ± 0.10 | 0.137 |
| BMI (kg/m<sup>2</sup>) | 25.54 ± 4.02 | 27.37 ± 5.15 | < 0.001* |
| FBG (mg/dL) | 91.76 ± 10.1 | 92.73 ± 15.4 | 0.891 |
| TC (mg/dL) | 152.3 ± 29.3 | 174.0 ± 32.8 | < 0.001* |
| LDL-C (mg/dL) | 87.89 ± 26.32 | 101.71 ± 28.43 | < 0.001* |
| HDL-C (mg/dL) | 50.72 ± 11.83 | 42.65 ± 10.63 | < 0.001* |
| Non -HDL Cholesterol (mg/dL) | 101.54 ± 27.57 | 131.34 ± 31.11 | < 0.001* |

SD: Standard Deviation; HTW-: Without diagnosis of Hypertriglyceridemic Waist; HTW+: With diagnosis of Hypertriglyceridemic Waist, PA Time: Physical Activity Time; BMI: Body Mass Index; FBG: Fasting Blood Glucose; TC: Total Cholesterol; HDL: Good Cholesterol (High Density Lipoproteins); LDL: Bad Cholesterol (Low Density Lipoproteins).
\* Significant values for $p \leq 0.05$

Table 2 shows the differences in the proportions between the total sample and the group diagnosed with HTW. Most adolescents diagnosed with HTW + (n = 125) were male, overweight and obese and with total cholesterol, HDL-C, LDL-C and non-HDL cholesterol.

**Table 2.**
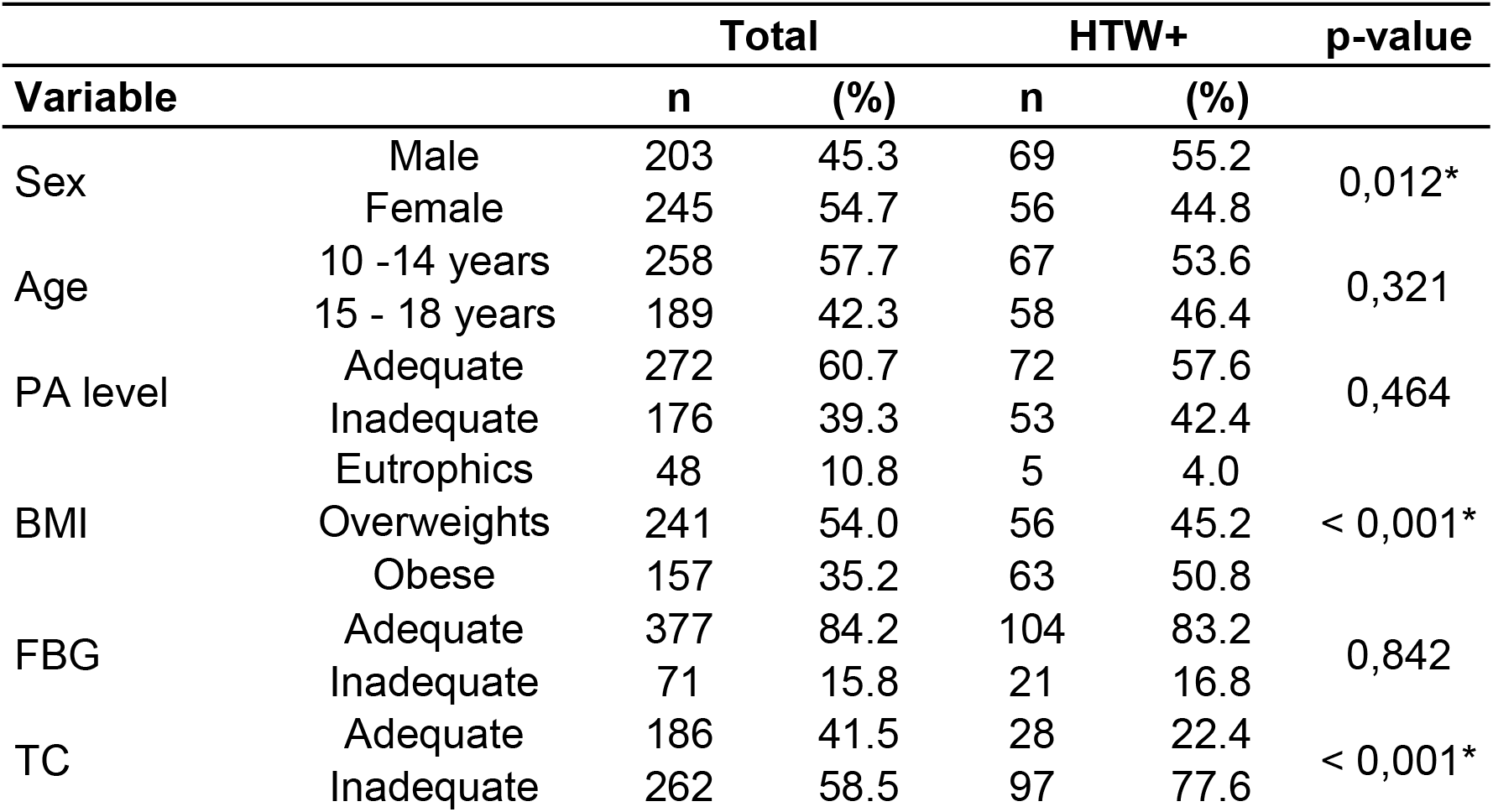
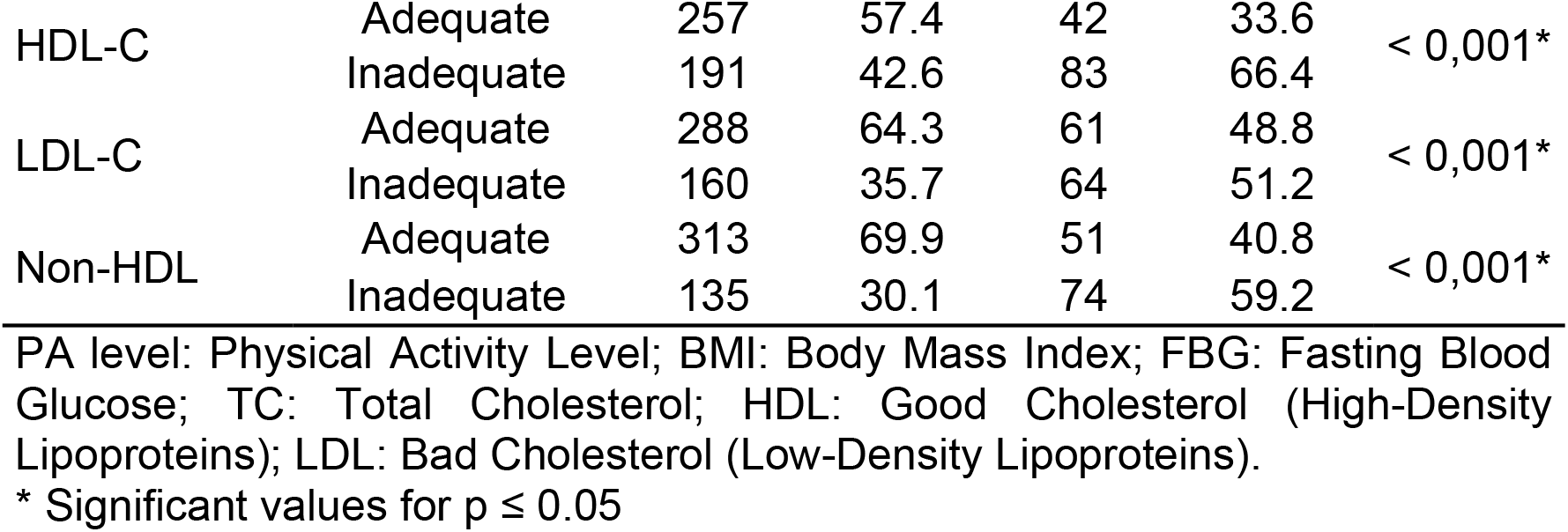
Physical activity time and health risk factors concerning HTW diagnoses in schoolchildren.

The age-adjusted logistic regression indicates that boys are 100% more 186 likely to be diagnosed with the HTW phenotype compared to girls. For nutritional status, overweight and obese schoolchildren are 140% and 520% more likely, respectively, compared to eutrophics. Finally, adolescents who had inadequate total cholesterol were 250% more likely to have the HTW phenotype (Table 3).

**Table 3.** Factors associated with HTW in schoolchildren from Paranavaí, measured by logistic regression.

| Variables |  | Adjusted OR<sup>#</sup> (IC<sub>95%</sub>) |
| --- | --- | --- |
| Sex | Female | 1 |
|  | Male | 2.0 (1.3 - 3.2)* |
| BMI | Eutrophics | 1 |
|  | Overweights | 2.4 (0.9 - 6.5) |
|  | Obese | 6.2 (2.7 - 17.2)* |
| TC | Adequate | 1 |
|  | Inadequate | 3.5 (2.1 - 5.7)* |

OR: Odds Ratios; BMI: Body Mass Index; TC: Total Cholesterol; #Adjusted by Age \* Significant Values for $p \leq 0.05$

## Discussion

The present study aimed to analyze the prevalence of the HTW phenotype and its associations with the level of physical activity and cardiovascular risk factors in students from public schools.

The findings of this research showed that among the 448 students evaluated, 125 had HTW, equivalent to 27.9% of the sample. Schoolchildren with HTW have an even higher body mass BMI, total cholesterol, LDL-C and non-HDL cholesterol and lower levels of HDL-C compared to the group not diagnosed with the phenotype. For the PA time, the groups did not present significant differences, however the average time, despite the great variation, was greater than the 300 minutes per week suggested for the age group ^13^ and according to the study by Alavian et al. (2008)^14^, low levels of physical activity increase the risk of adolescents with HTW.

Despite the lack of studies in the literature on the association of PA and HTW, the beneficial effects of increased physical activity in children and adolescents have been elucidated^15^, in different age groups, especially in variables related to body adiposity ^16,17^.This fact can be explained by the higher energy expenditure induced by the AF practice and the increased activity of hormones such as adrenaline and norepinephrine that have regulatory lipolytic action^18^.

In the case of the studied lipid variables (total cholesterol, HDL-C, LDL-C and non-HDL cholesterol), the present study demonstrated that individuals diagnosed with HTW have inadequate mean values in all variables. The study by Kumar et al., (2017)^19^, highlights that the detection of these variables in children and adolescents helps in the diagnosis of early dyslipidemia, a condition usually accompanied by obesity ^20^. This condition increases the appearance of atherosclerosis and cardiovascular diseases in adulthood ^21–24^.

Regarding fasting blood glucose, the values were similar between groups, which can be explained by the fact that this variable hardly changes in school ages ^25^. Normally, in this stage of life, obesity is related to the increase in circulating insulin levels (hyperinsulinemia), and it is only after the onset of insulin resistance that fasting blood glucose levels change, usually in adulthood.

As already mentioned, the HTW picture includes the combination of abdominal obesity and elevated TG (dyslipidemia) and, in the present study, 27.9% of students presented this diagnosis. In 2014, in the same population and city, with the same classification criteria, the prevalence of HTW was 20.7% ^6^, that is, in five years the incidence was 7.2%. Different prevalences have been reported in the literature, similar values have also been observed considering HTW assessments in different places around the world ^26,27^, and in Brazil ^2–28^, however, in all of them, the criteria and cutoff points were different.

Despite the differences in prevalence, there is consensus among the studies that the early diagnosis of HTW is of paramount importance, as it provides the identification of risk factors for the development of Metabolic Syndrome, which has a direct contribution to the onset of cardiovascular diseases ^29^. The symptomatic presence of only one of the variables is a sufficient parameter for specific interventions added to non-pharmacological strategies, highlighting the actions developed from the promotion of good life habits through the increase in daily PA practices in adolescence ^30^.

It is worth mentioning that the present research had limitations characteristic of a cross-sectional study regarding the causal relationships between the variables, as well as the absence of direct measures in the weekly PA time (unusual in epidemiological studies), although the present research used a questionnaire validated, the high standard deviation values in both groups may reflect overestimated or underestimated measures in the sample. Another limiting point of the research was the lack of analysis of circulating insulin levels, which would give better indications about glycemic responses in schoolchildren.

Thus, new studies comparing the time of PA in schoolchildren diagnosed and not diagnosed with HTW and who preferably use direct measures are necessary to elucidate whether, effectively, the time of PA differs between groups and whether this variable is associated or not with the phenotype.

In this sense, this research demonstrated real ecological applicability of the proposal, based on simple and clinically viable approaches for the identification of young people with HTW. As the present study was developed on a school basis, the importance of systematic interventions that can benefit this population is highlighted, given the increase in the percentage of individuals diagnosed with HTW in five years in the city. Despite the absence of an association between PA and HTW, the need to implement programs to encourage PA practice in schools is recommended, given its mitigating effect on cardiovascular risk factors in these populations.

## Acknowledgments

To the Paranavai Regional Education Center, to the principals of all public and school schools involved in the research

## References

1. Lemieux I, Pascot A, Couillard C, Lamarche B, Tchernof A, Alméras N, et al. Hypertriglyceridemic waist: A marker of the atherogenic metabolic triad (hyperinsulinemia; hyperapolipoprotein B; small, dense LDL) in men? Circulation. 2000.

2. Costa PR de F, Assis AMO, Cunha C de M, Pereira EM, Jesus G dos S de, Silva LEM da, et al. Hypertriglyceridemic Waist Phenotype and Changes in the Fasting Glycemia and Blood Pressure in Children and Adolescents Over One-Year Follow-Up Period. Arq Bras Cardiol [Internet]. 2017;1(1009):47–53.

3. Silva DR, Werneck AO, Collings PJ, Fernandes RA, Barbosa DS, Ronque ERV, et al. Physical activity maintenance and metabolic risk in adolescents. J Public Heal (United Kingdom). 2018.

4. Thivel D, Masurier J, Baquet G, Timmons BW, Pereira B, Berthoin S, et al. High-intensity interval training in overweight and obese children and adolescents: systematic review and meta-analysis. J Sports Med Phys Fitness [Internet]. 2018;59(2).

5. Buchan DS, Boddy LM, Despres JP, Grace FM, Sculthorpe N, Mahoney C, et al. Utility of the hypertriglyceridemic waist phenotype in the cardiometabolic risk assessment of youth stratified by body mass index. Pediatr Obes. 2016;11(4):292–8.

6. Guilherme FR, Molena-Fernandes CA, Hintze LJ, Fávero MTM, Cuman RKN, Rinaldi W. Hypertriglyceridemic Waist and Metabolic Abnormalities in Brazilian Schoolchildren. Medeiros R, editor. PLoS One [Internet]. 2014 Nov 14;9(11):e111724.

7. Conde WL, Monteiro CA. Body mass index cutoff points for evaluation of nutritional status in Brazilian children and adolescents. J Pediatr (Rio J) [Internet]. 2006 Aug 9;82(4):266–72.

8. Fernández J, Redden D, Pietrobelli A, Allison D. Waist circumference percentiles in nationally representative samples of African-American, European-American, and Mexican-American children and adolescents. J Pediatr. 2004;145:439–44.

9. Silva DR, Werneck AO, Collings PJ, Fernandes RA, Barbosa DS, Ronque ERV, et al. Physical activity maintenance and metabolic risk in adolescents. J Public Heal (United Kingdom). 2018;40(3):493–500.

10. Esmaillzadeh A, Mirmiran P, Azizi F. Clustering of metabolic abnormalities in adolescents with the hypertriglyceridemic waist phenotype. Am J Clin Nutr. 2006;83:36–46.

11. Farias Júnior JC de, Lopes A da S, Mota J, Santos MP, Ribeiro JC, Hallal PC. Validade e reprodutibilidade de um questionário para medida de atividade física em adolescentes: uma adaptação do Self-Administered Physical Activity Checklist. Rev Bras Epidemiol [Internet]. 2012 Mar;15(1):198–210.

12. Sallis JF, Strikmiller PK, Harsha DW, Feldman HA, Ehlinger S, Stone EJ, et al. Validation of interviewer- and self-administered physical activity checklists for fifth grade students. Med Sci Sport Exerc. 1996;28(7):840–51.

13. Strong WB, Malina RM, Blimkie CJR, Daniels SR, Dishman RK, Gutin B, et al. Evidence Based Physical Activity for School-age Youth. J Pediatr. 2005 Jun;146(6):732–7.

14. Alavian SM, Motlagh ME, Ardalan G, Motaghian M, Davarpanah AH, Kelishadi R. Hypertriglyceridemic waist phenotype and associated lifestyle factors in a National Population of Youths: CASPIAN study. J Trop Pediatr. 2008;54(3):169–77.

15. Barker AR, Gracia-Marco L, Ruiz JR, Castillo MJ, Aparicio-Ugarriza R, González-Gross M, et al. Physical activity, sedentary time, TV viewing, physical fitness and cardiovascular disease risk in adolescents: The HELENA study. Int J Cardiol [Internet]. 2018;254:303–9.

16. Sun S, Zhang H, Kong Z, Shi Q, Tong TK, Nie J. Twelve weeks of low volume sprint interval training improves cardio-metabolic health outcomes in overweight females. J Sports Sci [Internet]. 2019;37(11):1257–64.

17. Taylor JL, Holland DJ, Spathis JG, Beetham KS, Wisløff U, Keating SE, et al. Guidelines for the delivery and monitoring of high intensity interval training in clinical populations. Prog Cardiovasc Dis [Internet]. 2019;62(2):140–6.

18. Osiński W, Kantanista A. Physical activity in the therapy of overweight and obesity in children and adolescents. Needs and recommendations for intervention programs. Dev period Med. 2017;21(3):224–34.

19. Kumar S, Kelly AS. Review of Childhood Obesity: From Epidemiology, Etiology, and Comorbidities to Clinical Assessment and Treatment. Mayo Clin Proc [Internet]. 2017;92(2):251–65.

20. Nuotio J, Pitkänen N, Magnussen CG, Buscot MJ, Venäläinen MS, Elo LL, et al. Prediction of Adult Dyslipidemia Using Genetic and Childhood Clinical Risk Factors: The Cardiovascular Risk in Young Finns Study. Circ Cardiovasc Genet. 2017.

21. Webber LS, Srinivasan SR, Wattigney WA, Berenson GS. Tracking of serum lipids and lipoproteins from childhood to adulthood: The bogalusa heart study. Am J Epidemiol. 1991.

22. Li S, Chen W, Srinivasan SR, Bond MG, Tang R, Urbina EM, et al. Childhood Cardiovascular Risk Factors and Carotid Vascular Changes in Adulthood: The Bogalusa Heart Study. J Am Med Assoc. 2003;

23. Daniels SR, Greer FR. Lipid screening and cardiovascular health in childhood. Pediatrics. 2008.

24. Lozano P, Henrikson NB, Morrison CC, Dunn J, Nguyen M, Blasi PR, et al. Lipid screening in childhood and adolescence for detection of multifactorial dyslipidemia: Evidence report and systematic review for the US Preventive Services Task Force. JAMA - Journal of the American Medical Association. 2016.

25. Romualdo MCDS, De Nóbrega FJ, Escrivão MAMS. Insulin resistance in obese children and adolescents. J Pediatr (Rio J). 2014.

26. Kelishadi R, Jamshidi F, Qorbani M, Motlagh ME, Heshmat R, Ardalan G, et al. Association of hypertriglyceridemic-waist phenotype with liver enzymes and cardiometabolic risk factors in adolescents: the CASPIAN-III study. J Pediatr (Rio J). 2016;92(5):512–20.

27. Chen S, Guo X, Dong S, Yu S, Chen Y, Zhang N, et al. Association between the hypertriglyceridemic waist phenotype and hyperuricemia: a cross-sectional study. Clin Rheumatol. 2017.

28. Barreiro-Ribeiro F, Vasques ACJ, Da Silva CDC, Zambon MP, Rodrigues AMDB, Camilo DF, et al. Hypertriglyceridemic Waist Phenotype Indicates Insulin Resistance in Adolescents According to the Clamp Technique in the BRAMS Study. Child Obes. 2016.

29. Pereira PF, Faria FR De, Faria ER De, Hermsdorff HHM, Peluzio MDCG, Franceschini SDCC, et al. Indicadores antropométricos para identificar síndrome metabólica e fenótipo cintura hipertrigliceridêmica: uma comparação entre as três fases da adolescência. Rev Paul Pediatr. 2015.

30. Kuschnir MCC, Bloch KV, Szklo M, Klein CH, Barufaldi LA, De Azevedo Abreu G, et al. ERICA: Prevalence of metabolic syndrome in Brazilian adolescents. Rev Saude Publica. 2016;50(suppl 1):1s–13s.

